# End of preservation normothermic machine perfusion of porcine kidneys after ischaemic injury reprograms metabolism and induces fibrosis after transplant despite unchanged function: insights from the renal proteome

**DOI:** 10.1101/2024.09.27.615349

**Authors:** John Francis Mulvey, Corinna Snashall, Kaithlyn Rozenberg, M. Letizia Lo Faro, Marco Eijken, Stine Lohmann, Cyril Moers, Henri Leuvenink, Carla Baan, Martin Hoogduijn, Anna Krarup Keller, Chris Sutton, James Hunter, Bente Jespersen, Rutger Ploeg, Sadr Shaheed

## Abstract

Normothermic machine perfusion (NMP) after initial hypothermic preservation of donor kidneys prior to transplantation is becoming a clinical reality, but the precise molecular mechanisms through which the graft is impacted remain only partially characterised. Using an unbiased proteomic methodology, we found that auto transplantation of is chaemically injured porcine kidneys resulted in an activation of the stress response 14 days after transplantation, as well as in selective changes in the proteins responsible for the metabolism of organic acids. The addition of 4 hours of NMP at the end of organ preservation (endNMP) resulted in coordinated changes to the renal proteome at 14 days when compared with the effect of transplant after preservation by hypothermic machine perfusion alone: most notably increased fibrosis and widespread additional reprogramming of metabolism. These findings were supported by intersection with single cell transcriptomics data which suggested an enrichment of proteins predominantly expressed in fibroblasts in kidneys with end of preservation NMP 14 days post-transplant compared to healthy kidneys. Our data showed that the addition of endNMP to existing preservation strategies resulted in a different molecular phenotype after transplantation, despite unchanged filtration function. In addition to potentially conferring benefits, NMP may also result in potentially detrimental molecular changes and thus protocols should be carefully evaluated to derive optimal clinical outcomes.

## 1. Introduction

Preservation methods in organ transplantation are changing rapidly following decades where static cold storage of organs was the routine clinical practice. The causes of this change encompass a changing donor population^1^, an increased need to transplant marginal organs to meet the supply/demand mismatch that is causing long waitlists for patients requiring transplantation^2^, as well as the development of novel technology that means routine preservation by continuously perfusing organs is now clinically feasible. Such continuous perfusion has been well demonstrated to result in better outcomes for recipients when perfusion is conducted at hypothermic temperatures (4-12 degrees)^3^, but the organ remains far from normal physiology during the preservation period, with both a greatly reduced metabolic rate and injury induced by cold stress^4^. Oxygenation of the perfusate during hypothermic perfusion (oxHMP) aims to close this gap by supplying the graft with dissolved oxygen to support metabolism during preservation. This has been shown to result in partial recovery of high energy adenosine nucleotide levels^5,6^ as well as improvement in functional outcomes in preclinical models^7^. Clinically, it also results in better outcomes for recipients compared to static cold storage, at least in certain donor populations^8–10^. However, perfusion at normothermia may ultimately provide the greatest opportunity to recapitulate normal physiology during the preservation period, and by enabling a normal metabolic rate is likely to best support interventions to be performed upon the graft prior to transplantation that may be transformative in solid organ transplantation.

Normothermic machine perfusion (NMP) has been shown to be safe and feasible in the liver^11–15^, and also to result in decreased adverse outcomes: notably lower levels of graft injury and a lower organ discard rate. Since NMP for the entire preservation period presents significant logistical and economic hurdles, much research to date has focused on ‘end of preservation’ NMP (endNMP). This is commonly carried out for a duration of 1-4 hours following a conventional preservation method (such as static cold storage or hypothermic machine perfusion) for logistical reasons. Following success in the liver and positive evidence from preclinical models of kidney transplantation^16,17^, there are currently clinical trials underway or with recently published results to determine both safety and efficacy of NMP in the kidney^18–21^. Most recently, safety has been reported of endNMP up to 23 hours after prior cold storage in a clinical kidney feasibility trial followed by successful transplantation^18^. This paradigm is attractive since in addition to potential direct benefits of NMP, having isolated organs at normothermia can provide a platform to assess^22^, recondition and repair^23,24^ or genetically modify^25^ organs. To enable translation the proximal application of NMP is focused upon direct benefits to recipient outcomes with clinically relevant trial endpoints such as delayed graft function^21^ and one year graft function. However, the precise molecular mechanisms through which such a benefit may be conferred remain poorly elucidated in preclinical studies. Furthermore, it is also vital to characterise any detrimental effects that may emerge following NMP, particularly since the addition of NMP often results in an increase in the total preservation time and an increase in the time during which the graft is at risk of warm ischaemia.

We therefore focused our investigations upon the effects of NMP assessed 14 days after transplantation, in contrast to much of the existing preclinical literature which reports acute changes during and immediately following perfusion^26–31^. We utilised an unbiased proteomic methodology to assess the effects of preservation of kidneys by either oxygenated HMP alone or oxygenated HMP with the addition of 4 hours of NMP at the end of the preservation period. To best dissect the direct effects of NMP upon the graft, we did this in a porcine model of autotransplantation of kidneys after ischaemic injury^24^: without the transplant being exposed to a hostile immune environment or subject to immunosuppressive therapy.

## 2. Materials and Methods

### 2.1 Porcine autotransplantation model

We used a porcine model of autotransplantation after an injury period of 75 min warm ischaemia as previously reported^24^ and described in full in the supplementary content. These experiments were approved by the Danish Animal Experimentation Council (reference-number 2016-15-0201-01145) with local ethical approval. Healthy control kidneys were obtained from pigs that had been subjected to a non-invasive infusion of 25 mL phosphate buffered saline as a placebo procedure.

### 2.2. Sample collection and processing

At 14 days post-transplant or placebo procedure cortical punch biopsies (3 mm) were taken from the upper pole, medial, and lower pole locations of the kidney, placed in RNAlater (Thermo Fisher Scientific, UK) and stored at -80 °C before subsequent analysis. These 3 regions were subsequently combined to abrogate sampling artefacts. Details of protein extraction, digestion and peptide labelling are presented in the supplementary content. Healthy kidney tissue was obtained in the same way from pigs that had not been exposed to ischaemia and transplantation^32^.

### 2.3. LC-MS/MS analysis

Tandem mass tag (TMT) labelled peptides were pre-fractionated in order to reduce sample complexity using a 150 mm C18 column on an Agilent 1260 Infinity II high performance liquid chromatography instrument (Agilent Technologies, USA). Gradient elution was used to collect 24 fractions in 60 min, using a linear gradient of 2% to 35% acetonitrile at pH 9. These fractions were then agglomerated into 16 components as follows: 1, 2, 3, 4, 5, 6, 7, 8, 9, 10, 11, 12, 13-15, 16-18, 19-21, 22-24, and then dried under vacuum.

Online separation was performed using a Dionex Ultimate 3000-high performance liquid phase system (mobile phase A: 0.1% formic acid in water; mobile phase B: acetonitrile supplemented with 0.1% formic acid. Peptides were eluted with a 65-minute linear gradient: 2–25% solvent B (90% ACN, 0.1% FA), 10 minutes; 25–50% solvent B, 5 minutes; 50–90% solvent B, 8 minutes; column wash with 90% solvent B, followed by 16 minutes column equilibration with solvent A at a flow rate of 300 nL/min. Analysis was performed using a Orbitrap Fusion mass spectrometry equipped with a Nanoflex ion source (Thermo Fisher Scientific, UK). The parent ions of the peptide were detected and analysed using a high resolution orbitrap with ion source voltage 2.3 kV (MS^1^ scan range was set at 375–1500 m/z, with resolution of 120,000 at m/z 200. MS^2^ was performed using an ion-trap at rapid scan rate, with the following parameters: charge state 2-7, dynamic exclusion of 50 seconds, cycle time 3 seconds and collision induced dissociation ∼35%. Automated Synchronous Precursor Selection for MS^3^ was used for quantification of TMT10plex and performed on the orbitrap at 30,000 resolutions, with scan range 100-500 m/z, maximum injection time 105 ms and with 65% high-energy collisional dissociation. Methods to identify and quantify protein abundance are presented in the supplementary content.

### 2.4. Statistical analysis

Full details of methodology are described in the supplementary content. A false discovery rate of 5% using the method of Benjamini and Hochberg^33^ was considered significant. All data analyses and visualisation were performed using R^34^.

## 3. Results

### 3.1. Porcine Transplant

Ischaemically damaged kidneys were autotransplanted into pigs following either preservation by oxHMP alone or oxHMP + endNMP (hereafter referred to as endNMP) as previously reported^24^, or procured from pigs 14 days after receiving an infusion of phosphate buffered saline into the renal artery as a placebo procedure (healthy controls, Fig 1A). 14 days after either transplantation, glomerular filtration rate was measured using 99mTc-DTPA. GFR is known since the early 20^th^ century to be dependent upon kidney mass^35,36^. Kidney mass increased 14 days after transplantation in comparison to control but did not differ between groups (Fig 1B; F-test using mixed effects model, time point - *p* = 0.0001, preservation method – *p* = 0.08, interaction – *p* = 0.93). After normalising measured GFR to kidney mass in order to take into account the variation observed within experimental groups, no significant difference was found between kidneys transplanted following either oxHMP or oxHMP + endNMP, despite a long period of 75 min warm ischaemia (Fig 1C; Welch’s t-test, *p* = 0.55).

**Figure 1.**
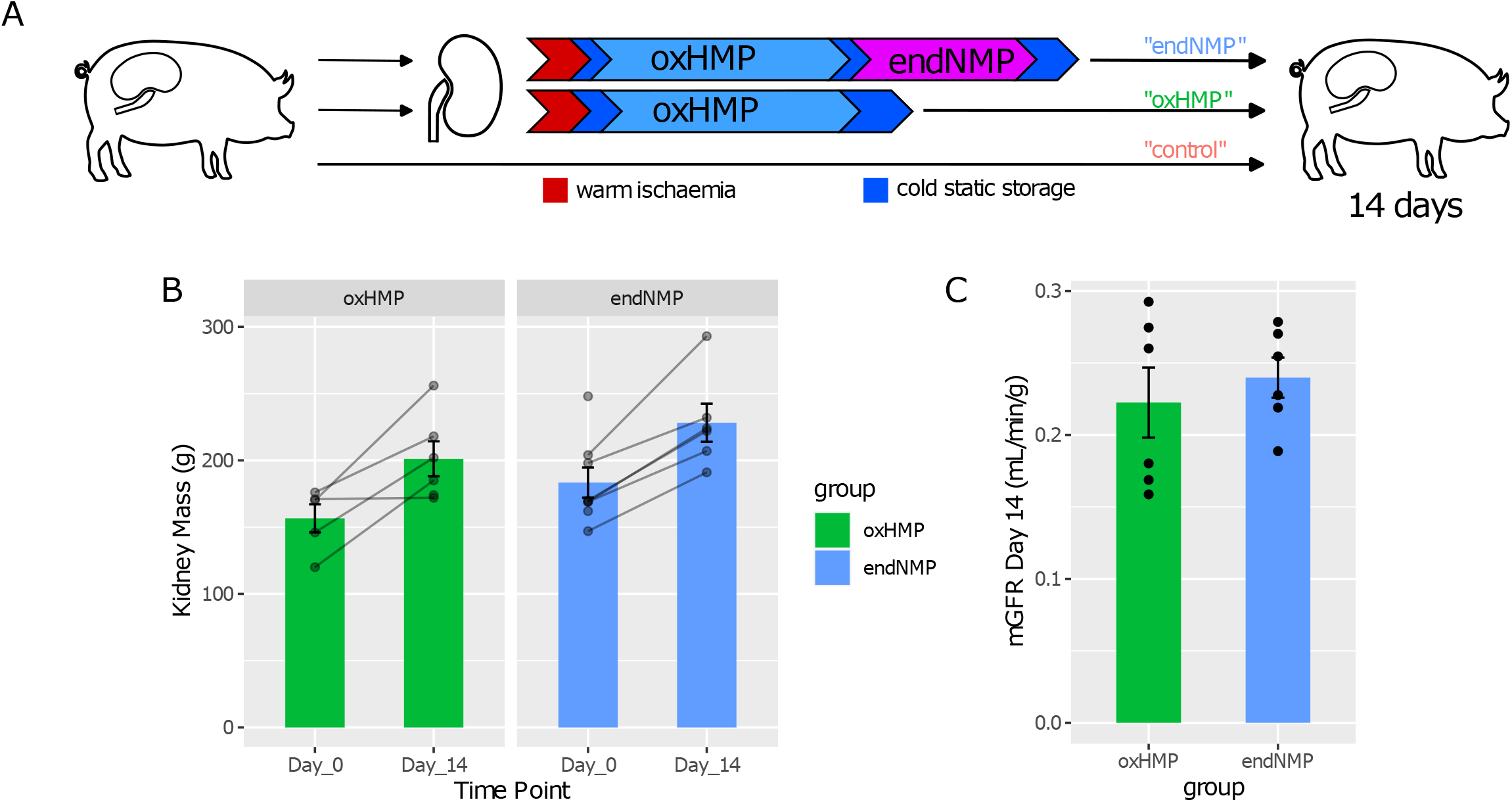
**A**. Left kidneys were subject to 75 min warm ischaemia and thenremoved from pigs (n=6-7). After a short period of cold static storage, they were preserved either by 14 hours oxygenated HMP alone, or this was followed by a further 4 hours of normothermic machine perfusion. Kidneys were then transplanted back into the pig of origin, and the contralateral kidney removed. Pigs were then monitored for 14 days before collection of tissue. Control kidneys were taken from pigs who similarly received 14 days of follow-up, after a placebo procedure. **B**. Kidneys increased in mass 14 days after transplantation (p = 0.0001), but this was not different between groups (preservation method – p = 0.08, interaction – p = 0.93). **C**. Measured glomerular filtration rate shows no significant difference between transplanted kidneys preserved by oxHMP alone or with endNMP when accounting for kidney mass (Welch’s t-test, p = 0.55). oxHMP - oxygenated hypothermic machine perfusion, endNMP – end of preservation normothermic machine perfusion, mGFR – measured glomerular filtration rate

### 3.2. Summary of proteomics

Biopsies were taken from kidneys 14 days after transplantation or the placebo procedure, proteins extracted and their abundances quantified by ultra-high resolution mass spectrometry after extensive fractionation (Fig 2A) resulting in 901 proteins quantified. Principal component analysis shows that the experimental groups correspond to the major sources of variation in the proteome profile (Fig 2B, Fig S1A), and accordingly there are 259 proteins whose expression differs statistically between control kidneys and preservation by oxHMP alone or by endNMP (Fig 2C-D, FDR < 0.05). There is a detectable signature of transplantation upon the renal proteome that remains 14 days later with 61 proteins statistically altered in the same direction in both oxHMP and endNMP groups, but surprisingly 74% (192/259) of the differentially regulated proteins in comparison to healthy control kidneys were specific to the mode of preservation (Fig 2D-E).

**Figure 2.**
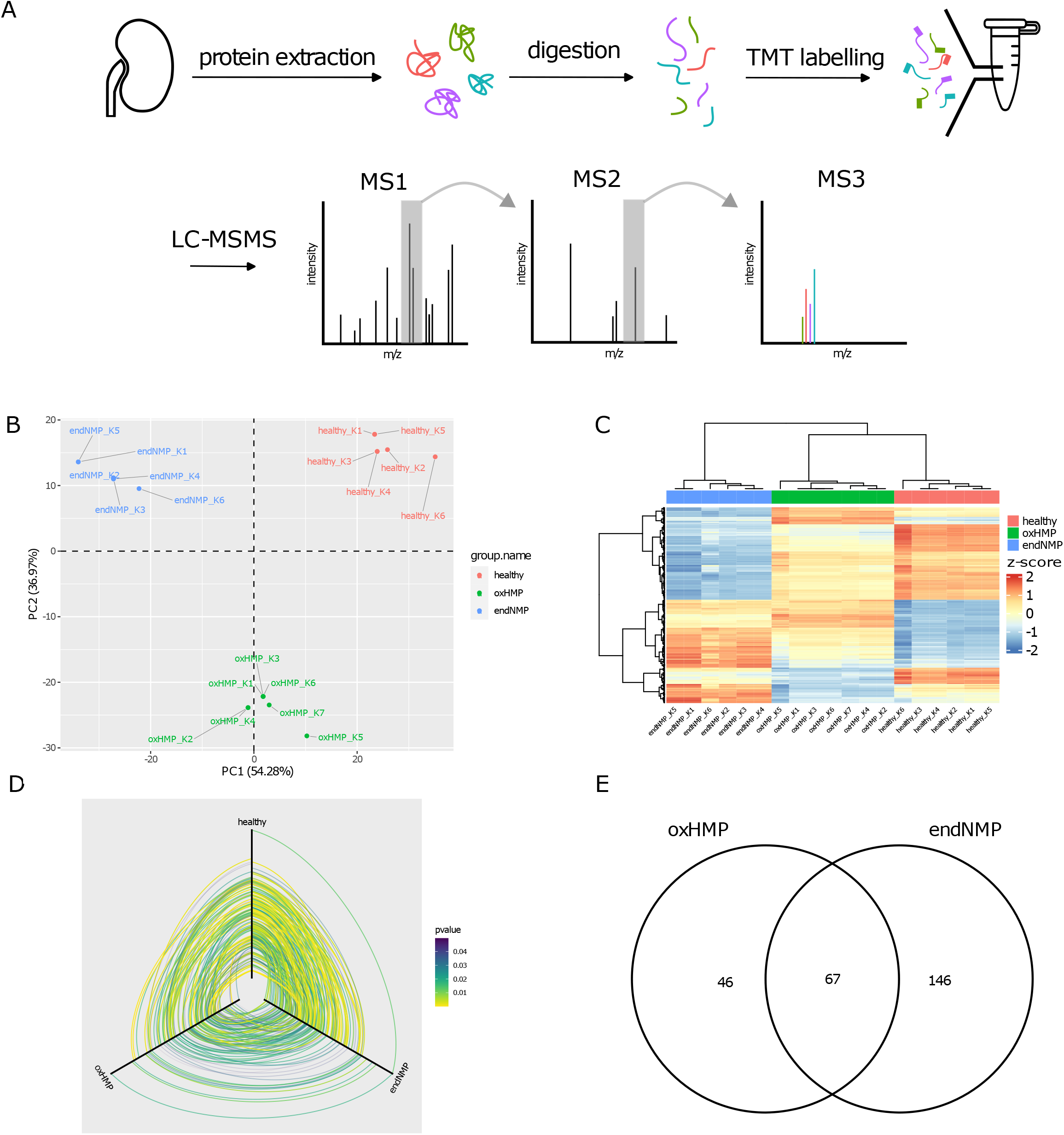
**A**. Biopsies taken from the upper, lateral and lower regions of each kidney were pooled, before proteins were extracted, digested into peptides and then labelled with TMT to allow multiplexing. Following 2-dimensional separation of peptides by liquid chromatography at both low and high pH, bottom-up proteomics was performed using mass spectrometry. At MS1, ions are selected for further analysis in MSn. Fragments are isolated in MS2 to ensure any co-eluting peptides are removed before quantification at MS3, where the relative abundance of the TMT labels in conjunction with peptide identifications from MS1 and MS2 provides accurate quantification of the relative abundances of proteins present in the original samples. **B**. Principal component analysis shows that the major sources in variation of protein expression separate the experimental groups. **C**. Heatmap of proteins significantly different between experimental groups, showing protein-wise z-scored abundance. **D**. Hive plot summarising the differences in protein expression. Each axis plots mean protein abundance in each condition. When a protein is significantly different between conditions, a line is plotted to indicate its mean abundance on the axis of the respective group with the colour indicating the strength of the statistical association. **E**. Comparison of the proteins that are significantly different from control kidneys in both the oxHMP and endNMP groups. There is a small overlap indicative of a long-lasting proteomic signature of transplantation, but the majority of differences seen are specific to the method of preservation. TMT – tandem mass tag, LC-MSMS – liquid chromatography tandem mass spectrometry

### 3.3. Effect of transplantation after preservation by oxHMP

The abundance of 113 proteins was significantly altered after transplantation with preservation by oxHMP in comparison to healthy kidneys, amongst which 66 proteins were upregulated and 48 proteins downregulated (Fig 3A). Enrichment of gene sets annotated as biological processes showed a sustained upregulation of stress response pathways, and a wide downregulation of proteins functioning in small molecule metabolic processes: particularly of organic acids including amino acids and carboxylic acids. The kidney is a tissue in which metabolism is highly segregated both in space and between cell types, and to understand this further we intersected our data with an atlas single cell transcriptomics dataset^37^. Using these data to define markers for particular cell types (Fig S1E), our proteomics data suggest a decrease in cells of the proximal tubule (Fig 3D, which we hypothesise to underlie this metabolic downregulation. Of the homologues of significantly downregulated genes after preservation by oxHMP, 46% (17 out of 37) are found to be markers of proximal tubule cells, and all are expressed in proximal tubule cells. These cells are highly metabolically active, and hence among the first segments of the kidney to be damaged by ischaemia.

**Figure 3.**
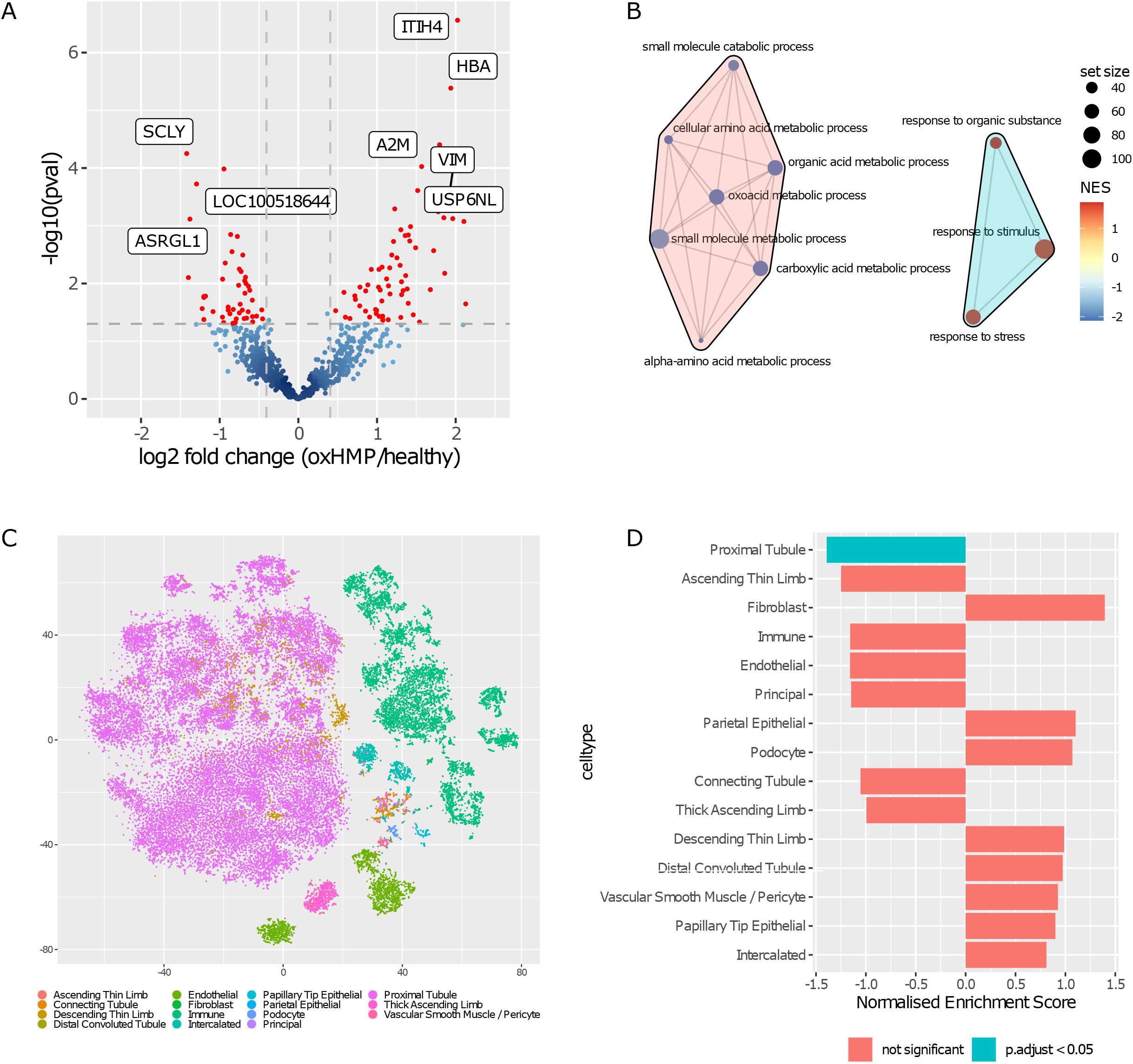
**A**. Proteins significantly different between transplanted kidneys preserved by oxHMP and healthy kidneys. **B**. Gene set enrichment analysis of gene ontology biological processes shows an upregulation of stress response pathways and a downregulation of metabolic processes in the metabolism of small organic acids remaining at 14 days after transplantation. **C**. t-distributed stochastic neighbour embedding summarising single cell transcriptomics of Stuart et al. (2019), according to cell type assigned from the transcriptome of each cell. **D**. Intersection with single cell transcriptomics indicates that there is a significant decrease in expression of proteins with specificity to proximal tubule cells after transplant. Normalised enrichment score quantifies the extent to which cell types are enriched within oxHMP or healthy kidneys. NES = normalised enrichment score

### 3.4. Effect of end of preservation NMP

The expression of 213 proteins were significantly altered 14 days after transplant with the addition of 4 hours endNMP following oxHMP in comparison to healthy kidney controls, amongst which 92 proteins were upregulated and 121 proteins downregulated (Fig 4A). Gene set enrichment analysis suggested a much more widespread downregulation of metabolism in comparison to oxHMP alone. This again encompassed the metabolism of small organic acids but in addition of hexose and lipid metabolism: together comprising the three major sources of fuel for respiration (Fig 4B). Seven gene sets were commonly downregulated in both oxHMP and oxHMP + endNMP in comparison to healthy kidney controls, all related to the metabolism of organic acids such as amino acids and carboxylic acids (Fig S1B).

**Figure 4.**
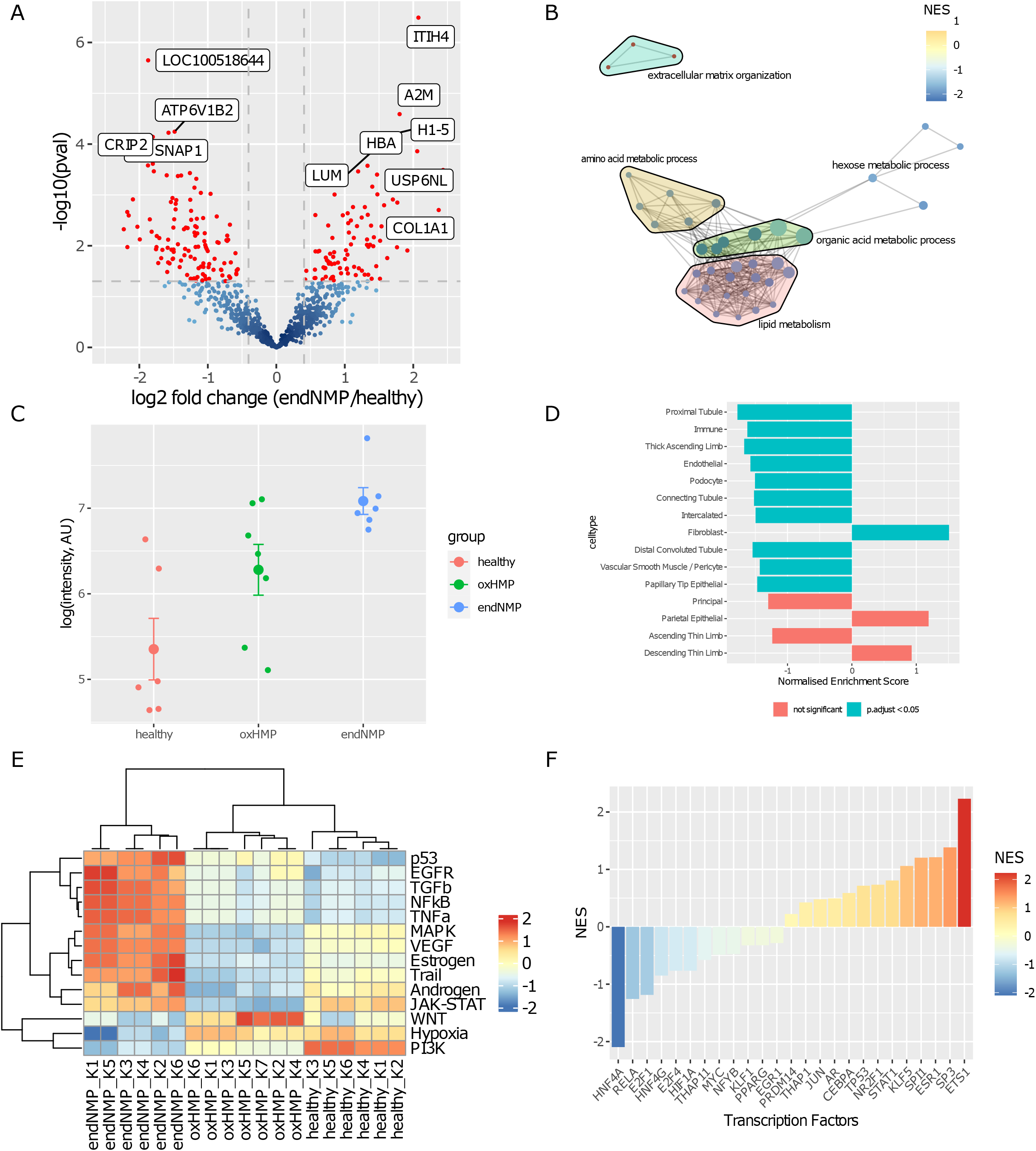
**A**. Proteins significantly different between transplanted kidneys preserved with the addition of 4 hours end of preservation NMP and healthy kindeys. **B**. Gene set enrichment analysis of gene ontology biological processes. **C**. Expression of COL1A1 measured by western blotting, normalised to β-actin. **D**. Intersection with single cell transcriptomics indicates that there is an increase in proteins originating from fibroblasts, whilst a negative enrichment of proteins from the majority of other cell types of the kidney. **E**. Calculating activities of common cellular signalling pathways shows alterations in, for example, hypoxia, MAPK, EGFR, TGFb and TNFa signalling activity. **F**. Functional activity of transcription factors between endNMP and healthy kidneys, calculated based on the expression of regulated proteins. Activity of, for example, HIF1A and HNF4A is decreased whilst that of ETS1 is increased. NES – normalised enrichment score, MAPK - mitogen-activated protein kinase, EGFR – epidermal growth factor receptor, TGFb–transforming growth factor beta, TNFa – tumour necrosis factor alpha, data show mean ± standard error where applicable

In kidneys transplanted after preservation by endNMP there is in addition an enrichment of processes related to organisation of the extracellular matrix in comparison to healthy kidneys which was not present in the oxHMP alone group, and is suggestive of fibrosis. COL1A1 is used as a canonical marker of fibrosis and was significantly upregulated in endNMP in our LC-MSMS proteomics (fold change in endNMP vs control = 5.2, *p* = 0.0002, Fig S1D). We also used western blotting to confirm the results by an orthogonal, albeit semiquantitative, method. This confirmed significantly higher expression of COL1A1 (*p* = 0.003) in the endNMP group compared to the healthy control group (Fig 4C, Fig S2B). Congruent with this, intersection with single cell transcriptomic profiles suggested an increase in fibroblasts (Fig 4D). Together with an increase in extracellular matrix proteins, this may explain the negative enrichment scores seen in most other common cell types of the kidney (Fig 4D), which are indicative of decreased abundance of these cell types.

We sought to understand the reasons behind this increase in fibrosis, and looked for evidence of molecular pathways whose activation is suggested by the proteomic profile. This indicated activation of TGFβ, TNFα and epidermal growth factor receptor (EGFR) pathways in the endNMP group in comparison to both oxHMP and healthy kidneys, which implies an increase in fibrosis (Fig 4E). Furthermore there was evidence for a decrease in the activity of the hypoxia pathway specifically in endNMP, whilst MAPK signalling was increased by oxHMP + endNMP whilst decreased by oxHMP alone. We further sought to determine if there were transcription factors whose changes in activity could be responsible for the observed changes in protein abundance. We observed functional evidence for an increase in the activity of the transcription factor protein C-ets-1 (ETS1) and a decrease in the activity of hepatocyte nuclear factor 4-alpha (HNF4A) in endNMP compared to healthy control kidneys (Fig 4F).

## 4. Discussion

In this study we examined the impact of endNMP upon the renal proteome 14 days after transplantation in a porcine model of autotransplantation, comparing this preservation strategy to kidneys perfused by oxHMP alone in the context of the proteome of healthy control kidneys. Unbiased proteomic methods revealed that endNMP induced a fibrotic signature compared to healthy kidneys with suggestions of a decrease in cells from a number of regions of the nephron. Much of the current preclinical literature has focused on either the effects of NMP alone^26,38^, followed by an *ex vivo* model of reperfusion^27,39^ or acute changes occurring after transplantation^40^. We believe that examining the effects 14 days after transplantation increases the relevance to clinical outcomes beyond primary graft function, which are known to be better predictors of long term graft survival^41^.

### 4.1. Response to transplantation

Hypothermic Machine Perfusion (HMP) is a current gold standard method by which to preserve kidneys *ex vivo* before transplantation, with current clinical evidence showing better recipient outcomes than cold static storage^8,10^. Supplementary administration of oxygen may result in better function and reduce rejection rates compared to non-oxygenated hypothermic machine perfusion in at least a subset of deceased donors^9,42^. The alterations in the proteome observed following oxHMP in our study are therefore relevant to the current clinical scenario, although titrated for an injurious period of warm ischaemia to mimic the scenario of donation after circulatory death. Our data show that the upregulation of stress response pathways is maintained 14 days after transplantation following oxHMP, but this was not observed following endNMP despite a 4-hour increase of the preservation period.

### 4.2. Metabolism

After transplantation following oxHMP there was a downregulation of carboxylic acid related metabolic processes, such as occur within and adjacent to the TCA cycle, when compared to healthy control kidneys. Metabolic regulation occurs not only through changes in the concentration of the enzymes responsible, but also through both processes such as post-translational modifications, allosteric regulation and the concentration of both substrates and products. It is likely that the extent of metabolic reprogramming will be underestimated by the changes in protein abundance alone. The addition of a short period of end of preservation NMP induced a much more widespread metabolic reprogramming when compared to the effect of transplantation after preservation by oxHMP alone. There was in addition a downregulation of amino acid, lipid and hexose metabolism: together comprising the three major fuel sources for respiration. The decrease in the metabolism of carboxylic acids affects mainly the tricarboxylic acid cycle. We observed for instance a decrease in the expression of isocitrate dehydrogenase 1 (IDH1), which is consistent with recent findings from metabolic flux in clinical kidney ischaemia reperfusion injury that suggest that the conversion of D-isocitrate to α-ketoglutarate is reduced^43^.

However, our intersection with single cell transcriptomics suggested that a decrease in proximal tubule cells also contributed to this difference, with decreased abundance of proteins from the proximal tubule cells observed in both oxHMP alone and endNMP compared to control kidneys. Proximal tubule cells have a high metabolic rate in order to meet the energetic demands required for the reabsorption of the bulk of sodium and other solutes, including glucose and amino acids, that occurs in this segment of the nephron. The “inner stripe” containing much of the proximal tubule cell mass is the most susceptible region of the kidney to ischaemic injury^44^, since it operates at the intersection between an already low oxygen tension during normal physiology^45^ and high energetic demands.

#### 4.2.1. Upstream regulators

We interrogated our data in combination with literature datasets of proteins altered by common signalling pathways, which predicted a decrease in hypoxia signalling pathways in endNMP in comparison to not only oxHMP alone but also control kidneys. This was corroborated by the abundances of proteins regulated by the transcription factor HIF1A, which decreased after endNMP in comparison to the control group. HIF1A is constitutively active at low levels in most cells, and we speculate that this dysregulation after endNMP may have contributed to the altered metabolic phenotype that we observed.

Our data on the abundance of proteins involved in metabolism measured at 14 days post-transplant stand in contrast to previous studies of the acute response to NMP. McEvoy and colleagues similarly employed an autotransplantation model comparing preservation with 8 hours of continuous NMP followed by 3 days follow-up, after which they performed proteomics on tissue biopsies. At this acute stage they observe an upregulation of fatty acid ß-oxidation, and the TCA cylcle^40^, directly opposes to our observations at 14 days. In human kidneys that were procured and subsequently discarded, 24 hours of NMP without transplantation has been reported to upregulate proteins involved in the TCA cycle^38^; given our data, NMP alone appears to produce a different metabolic phenotype in comparison to that which we observed 14 days after autotransplantation.

### 4.3. Fibrosis

A major hallmark of the phenotype of endNMP - but not oxHMP alone - in our data was fibrosis, which is well described following ischaemic injury to the extent that ischaemia is commonly used to create animal models of renal fibrosis^46^. We observed upregulation of extracellular matrix proteins after endNMP compared to healthy control kidneys. The renal extracellular matrix is a complex network of fibrillar collagens, elastin, and various glycoproteins, forming for example basement membranes and the interstitial space. This is crucial to maintain tissue structure and function in the kidney, but in fibrosis the extracellular matrix replaces parenchymal tissue to the detriment of organ function. As an exemplar, the abundance of fibrillar collagen type 1 A1 (COL1A1) was significantly increased in the endNMP group compared to controls but there was no significant difference between the oxHMP alone and healthy groups. We have previously reported no difference in COL1A1 mRNA expression between oxHMP and endNMP groups^24^, with the expression levels in both elevated in comparison to healthy control kidneys. The poor correlation between transcript and protein abundance has been widely described^47,48^, and we argue this emphasises the value in measuring protein levels directly. In contrast, our previous histological findings showed a greater average histological score for fibrosis biopsies preserved by endNMP than oxHMP alone, although this was not statistically significant^24^. Our analysis here also suggested that endNMP corresponds to an increase in fibroblasts and a decrease in the majority of other cell types compared to healthy kidneys, corroborating the fibrotic phenotype.

#### 4.3.1. Upstream regulators

Our data suggested that NMP also induces an increase in the MAPK pathway, further than preservation by oxHMP alone. MAPK has been described to cause fibrosis across a number of physiological systems^49–52^, but has specifically been implicated in the pathogenesis of acute kidney injury, including that initiated by hypoxia^53–55^ where its activation has been reported to lead to an increase in apoptosis^56^. We also observed functional evidence for an increase in the activity of the transcription factors ETS1 after preservation by both oxHMP alone and endNMP, which has been described to result in remodelling of the extracellular matrix through increase in the secretion of matrix metalloproteases^57,58^.

### 4.4. Clinical relevance

endNMP may have advantages over hypothermic preservation methods alone in a number of respects^59^, but during its implementation care must be taken in order to mitigate any potential negative effects. Whilst there is a tendency to view normothermic machine perfusion as a single intervention, there are a range of different protocols that employ differing methods to keep the kidney in a quasi-physiological state *ex vivo*. Due to the increase in temperature and consequently metabolic rate in comparison to conventional preservation at hypothermia, there is an increased risk of hypoxic damage to the kidney, as suggested by the proteomic phenotype that we observed here. The most notable opportunities for warm ischaemia occur when starting or ending normothermic perfusion, if a mismatch should occur between the metabolic demands and the quantity of oxygen delivered by the perfusate. Strategies to manage the transition between hypothermia and normothermia are already being investigated^60^. It is also of note that the kidney must not only be supplied with oxygenated perfusate but must also uptake the oxygen from it, for which the delivery of oxygen in the perfusate is necessary but not sufficient^61^. That this is occurring can be validated in a closed system by measuring arterio-venous oxygen differences across the kidney.

### 4.5. Limitations

Research suggests that tissue from older and comorbid animals is more susceptible to ischaemia than that from young and healthy animals^62,63^, which is one of the key differences between the clinical population of organ donors and recipients and the young and otherwise healthy animals used here. We expect that our model therefore likely understates the importance of any period of warm ischaemia that occurs during endNMP.

We consciously used autotransplantation as a model system in order to dissect the differences in comparison to the clinical scenario where the allogenic immune response may have additional interactive effects with the mode of preservation. Our experimental design does not capture interactions between the immune response induced by allogeneic tissue types and the preservation method. Whilst the pig model enables these experiments to be performed ethically, there are also clearly differences between porcine and human physiology.

### 4.6. Conclusions

The addition of endNMP during the preservation period did not to result in differences in kidney function when compared to perfusion by oxHMP alone despite the increase in the total preservation time: our data showed no difference in glomerular filtration rate in kidneys transplanted after preservation by oxHMP alone or with the addition of endNMP. However, unbiased proteomic analysis suggests that adding endNMP after oxHMP induced a molecular phenotype of increased fibrosis and altered metabolic function. Protocols for endNMP should be carefully designed in order to minimise the insults that may be responsible for detrimental changes in addition to maximising benefits.

## Acknowledgements

We would like to thank all those involved in the care of the pigs in the animal house throughout the experimental procedures.

The study was funded by the Lundbeck Foundation.

## Disclosures

The authors of this manuscript have no conflicts of interest to disclose.

## Supporting information

Additional supporting information may be found online in the Supporting Information section.

## Data availability statement

The mass spectrometry proteomics data have been deposited to the ProteomeXchange Consortium via the PRIDE repository with the dataset identifier PXD048983. All code to reproduce our analysis will be made available following publication at 10.5281/zenodo.13146295.

## Abbreviations

DTPA: diethylenetriaminepentaacetic acid
EGFR: epidermal growth factor receptor
endNMP: end of preservation normothermic machine perfusion
ETS1: Protein C-ets-1
GFR: glomerular filtration rate
HNF4A: hepatocyte nuclear factor 4-alpha
mGFR: measured glomerular filtration rate
NMP: normothermic machine perfusion
oxHMP: oxygenated hypothermic machine perfusion
SDS-PAGE: sodium dodecyl sulphate-polyacrylamide gel electrophoresis
TMT: tandem mass tags
UW-CSS: University of Wisconsin cold storage solution

## Notes

### Competing Interest Statement

The authors have declared no competing interest.

## References

1. Bailey, P.K., Wong, K., Robb, M., Burnapp, L., Rogers, A., Courtney, A., and Wroe, C. (2020). Has the UK living kidney donor population changed over time? A cross-sectional descriptive analysis of the UK living donor registry between 2006 and 2017. BMJ Open 10, e033906. 10.1136/bmjopen-2019-033906.

2. NHS Blood and Transplant, N.E. (2022). Annual Report on Kindey Transplantation.

3. Taylor, M.J., and Baicu, S. C. (2010). Current State of Hypothermic Machine Perfusion Preservation of Organs: The Clinical Perspective. Cryobiology 60, S20–S35. 10.1016/j.cryobiol.2009.10.006.

4. Belzer, F.O., and Southard, J.H. (1988). Principles of solid-organ preservation by cold storage. Transplantation 45, 673–676.

5. Ravaioli, M., Baldassare, M., Vasuri, F., Pasquinelli, G., Laggetta, M., Valente, S., Pace, V.D., Neri, F., Siniscalchi, A., Zanfi, C., et al. (2018). Strategies to Restore Adenosine Triphosphate (ATP) Level After More than 20 Hours of Cold Ischemia Time in Human Marginal Kidney Grafts. Ann. Transplant. 23, 34–44. 10.12659/AOT.905406.

6. Darius, T., Vergauwen, M., Smith, T., Gerin, I., Joris, V., Mueller, M., Aydin, S., Muller, X., Schlegel, A., Nath, J., et al. (2020). Brief O2 uploading during continuous hypothermic machine perfusion is simple yet effective oxygenation method to improve initial kidney function in a porcine autotransplant model. Am. J. Transplant. 20, 2030–2043. 10.1111/ajt.15800.

7. Thuillier, R., Allain, G., Celhay, O., Hebrard, W., Barrou, B., Badet, L., Leuvenink, H., and Hauet, T. (2013). Benefits of active oxygenation during hypothermic machine perfusion of kidneys in a preclinical model of deceased after cardiac death donors. J. Surg. Res. 184, 1174–1181. 10.1016/j.jss.2013.04.071.

8. Moers, C., Smits, J.M., Maathuis, M.-H.J., Treckmann, J., van Gelder, F., Napieralski, B.P., van Kasterop-Kutz, M., van der Heide, J.J.H., Squifflet, J.-P., van Heurn, E., et al. (2009). Machine perfusion or cold storage in deceased-donor kidney transplantation. N. Engl. J. Med. 360, 7–19. 10.1056/NEJMoa0802289.

9. Jochmans, I., Brat, A., Davies, L., Hofker, H.S., van de Leemkolk, F.E.M., Leuvenink, H.G.D., Knight, S.R., Pirenne, J., Ploeg, R.J., and COMPARE Trial Collaboration and Consortium for Organ Preservation in Europe (COPE) (2020). Oxygenated versus standard cold perfusion preservation in kidney transplantation (COMPARE): a randomised, double-blind, paired, phase 3 trial. Lancet Lond. Engl. 396, 1653–1662. Leeuwen.

10. Jochmans, I., Moers, C., Smits, J.M., Leuvenink, H.G.D., Treckmann, J., Paul, A., Rahmel, A., Squifflet, J.-P., van Heurn, E., Monbaliu, D., et al. (2010). Machine perfusion versus cold storage for the preservation of kidneys donated after cardiac death: a multicenter, randomized, controlled trial. Ann. Surg. 252, 756–764. 10.1097/SLA.0b013e3181ffc256.

11. Nasralla, D., Coussios, C.C., Mergental, H., Akhtar, M.Z., Butler, A.J., Ceresa, C.D.L., Chiocchia, V., Dutton, S.J., García-Valdecasas, J.C., Heaton, N., et al. (2018). A randomized trial of normothermic preservation in liver transplantation. Nature 557, 50– 56. 10.1038/s41586-018-0047-9.

12. Markmann, J.F., Abouljoud, M.S., Ghobrial, R.M., Bhati, C.S., Pelletier, S.J., Lu, A.D., Ottmann, S., Klair, T., Eymard, C., Roll, G.R., et al. (2022). Impact of Portable Normothermic Blood-Based Machine Perfusion on Outcomes of Liver Transplant: The OCS Liver PROTECT Randomized Clinical Trial. JAMA Surg. 157, 189–198. 10.1001/jamasurg.2021.6781.

13. Ravikumar, R., Jassem, W., Mergental, H., Heaton, N., Mirza, D., Perera, M.T.P.R., Quaglia, A., Holroyd, D., Vogel, T., Coussios, C.C., et al. (2016). Liver Transplantation After Ex Vivo Normothermic Machine Preservation: A Phase 1 (First-in-Man) Clinical Trial. Am. J. Transplant. 16, 1779–1787. 10.1111/ajt.13708.

14. Ceresa, C.D.L., Nasralla, D., Watson, C.J.E., Butler, A.J., Coussios, C.C., Crick, K., Hodson, L., Imber, C., Jassem, W., Knight, S.R., et al. (2019). Transient Cold Storage Prior to Normothermic Liver Perfusion May Facilitate Adoption of a Novel Technology. Liver Transplant. Off. Publ. Am. Assoc. Study Liver Dis. Int. Liver Transplant. Soc. 25, 1503–1513. 10.1002/lt.25584.

15. Ghinolfi, D., Rreka, E., De Tata, V., Franzini, M., Pezzati, D., Fierabracci, V., Masini, M., Cacciatoinsilla, A., Bindi, M.L., Marselli, L., et al. (2019). Pilot, Open, Randomized, Prospective Trial for Normothermic Machine Perfusion Evaluation in Liver Transplantation From Older Donors. Liver Transpl. 25, 436. 10.1002/lt.25362.

16. Mazilescu, L.I., Urbanellis, P., Kaths, M.J., Ganesh, S., Goto, T., Noguchi, Y., John, R., Konvalinka, A., Mucsi, I., Ghanekar, A., et al. (2021). Prolonged Normothermic Ex Vivo Kidney Perfusion Is Superior to Cold Nonoxygenated and Oxygenated Machine Perfusion for the Preservation of DCD Porcine Kidney Grafts. Transplant. Direct 7, e751. 10.1097/TXD.0000000000001218.

17. Mazilescu, L.I., Goto, T., John, R., Rosales, R., Ganesh, S., Yu, F., Noguchi, Y., Kawamura, M., Dezard, V., Gao, F., et al. (2024). Combining Oxygenated Cold Perfusion With Normothermic Ex Vivo Perfusion Improves the Outcome of Donation After Circulatory Death Porcine Kidney Transplantation. Transplantation 108, 184–191. 10.1097/TP.0000000000004734.

18. ISRCTN - ISRCTN13292277: Investigation of the safety and feasibility of preservation of kidneys for up to 24 hours at normal body temperature prior to transplant 10.1186/ISRCTN13292277.

19. ISRCTN - ISRCTN91315246: Quality assessment of kidneys by ex-vivo warm perfusion prior to transplantation 10.1186/ISRCTN91315246.

20. Rijkse, E., Bouari, S., Kimenai, H.J.A.N., de Jonge, J., de Bruin, R.W.F., Slagter, J.S., van den Hoogen, M.W.F., IJzermans, J.N.M., Hoogduijn, M.J., and Minnee, R.C. (2021). Additional Normothermic Machine Perfusion Versus Hypothermic Machine Perfusion in Suboptimal Donor Kidney Transplantation: Protocol of a Randomized, Controlled, Open-Label Trial. Int. J. Surg. Protoc. 25, 227–237. 10.29337/ijsp.165.

21. Hosgood, S.A., Callaghan, C.J., Wilson, C.H., Smith, L., Mullings, J., Mehew, J., Oniscu, G.C., Phillips, B.L., Bates, L., and Nicholson, M.L. (2023). Normothermic machine perfusion versus static cold storage in donation after circulatory death kidney transplantation: a randomized controlled trial. Nat. Med. 29, 1511–1519. 10.1038/s41591-023-02376-7.

22. Hosgood, S.A., Thompson, E., Moore, T., Wilson, C.H., and Nicholson, M.L. (2018). Normothermic machine perfusion for the assessment and transplantation of declined human kidneys from donation after circulatory death donors. Br. J. Surg. 105, 388–394. 10.1002/bjs.10733.

23. Pool, M., Eertman, T., Sierra Parraga, J., ‘t Hart, N., Roemeling-van Rhijn, M., Eijken, M., Jespersen, B., Reinders, M., Hoogduijn, M., Ploeg, R., et al. (2019). Infusing Mesenchymal Stromal Cells into Porcine Kidneys during Normothermic Machine Perfusion: Intact MSCs Can Be Traced and Localised to Glomeruli. Int. J. Mol. Sci. 20, 3607. 10.3390/ijms20143607.

24. Lohmann, S., Pool, M.B.F., Rozenberg, K.M., Keller, A.K., Moers, C., Møldrup, U., Møller, B.K., Lignell, S.J.M., Krag, S., Sierra-Parraga, J.M., et al. (2021). Mesenchymal stromal cell treatment of donor kidneys during ex vivo normothermic machine perfusion: A porcine renal autotransplantation study. Am. J. Transplant. 21, 2348–2359. 10.1111/ajt.16473.

25. Yuzefovych, Y., Valdivia, E., Rong, S., Hack, F., Rother, T., Schmitz, J., Bräsen, J.H., Wedekind, D., Moers, C., Wenzel, N., et al. (2020). Genetic Engineering of the Kidney to Permanently Silence MHC Transcripts During ex vivo Organ Perfusion. Front. Immunol. 11, 265. 10.3389/fimmu.2020.00265.

26. Zaza, G., Neri, F., Bruschi, M., Granata, S., Petretto, A., Bartolucci, M., di Bella, C., Candiano, G., Stallone, G., Gesualdo, L., et al. (2023). Proteomics reveals specific biological changes induced by the normothermic machine perfusion of donor kidneys with a significant up-regulation of Latexin. Sci. Rep. 13, 5920. 10.1038/s41598-023-33194-z.

27. Hunter, J.P., Faro, L.L., Rozenberg, K., Dengu, F., Ogbemudia, A., Weissenbacher, A., Mulvey, J.F., Knijff, L., Gopalakrishnan, K., and Ploeg, R.J. (2022). Assessment of Mitochondrial Function and Oxygen Consumption Measured During Ex Vivo Normothermic Machine Perfusion of Injured Pig Kidneys Helps to Monitor Organ Viability. Transpl. Int. 0. 10.3389/ti.2022.10420.

28. Tietjen, G.T., Hosgood, S.A., DiRito, J., Cui, J., Deep, D., Song, E., Kraehling, J.R., Piotrowski-Daspit, A.S., Kirkiles-Smith, N.C., Al-Lamki, R., et al. (2017). Nanoparticle targeting to the endothelium during normothermic machine perfusion of human kidneys. Sci. Transl. Med. 9, eaam6764. 10.1126/scitranslmed.aam6764.

29. DiRito, J.R., Hosgood, S.A., Reschke, M., Albert, C., Bracaglia, L.G., Ferdinand, J.R., Stewart, B.J., Edwards, C.M., Vaish, A.G., Thiru, S., et al. (2021). Lysis of cold-storage-induced microvascular obstructions for ex vivo revitalization of marginal human kidneys. Am. J. Transplant. 21, 161–173. 10.1111/ajt.16148.

30. Blum, M.F., Liu, Q., Soliman, B., Dreher, P., Okamoto, T., Poggio, E.D., Goldfarb, D.A., Baldwin, W.M., and Quintini, C. (2017). Comparison of normothermic and hypothermic perfusion in porcine kidneys donated after cardiac death. J. Surg. Res. 216, 35–45. 10.1016/j.jss.2017.04.008.

31. Hosgood, S.A., Patel, M., and Nicholson, M.L. (2013). The conditioning effect of ex vivo normothermic perfusion in an experimental kidney model. J. Surg. Res. 182, 153–160. 10.1016/j.jss.2012.08.001.

32. Munk, A., Duvald, C.S., Pedersen, M., Lohmann, S., Keller, A.K., Møller, B.K., Ringgaard, S., Buus, N.H., Jespersen, B., and Eijken, M. (2022). Dosing Limitation for Intra-Renal Arterial Infusion of Mesenchymal Stromal Cells. Int. J. Mol. Sci. 23, 8268. 10.3390/ijms23158268.

33. Benjamini, Y., and Hochberg, Y. (1995). Controlling the False Discovery Rate: A Practical and Powerful Approach to Multiple Testing. J. R. Stat. Soc. Ser. B Methodol. 57, 289–300.

34. R Core Team (2022). R: A Language and Environment for Statistical Computing. (R Foundation for Statistical Computing).

35. Taylor, F.B., Drury, D.R., and Addis, T. (1923). The regulation of renal activity. Am. J. Physiol.-Leg. Content 65, 55–61. 10.1152/ajplegacy.1923.65.1.55.

36. Addis, T., Shevky, A.E., and Bevier, G. (1918). The regulation of renal activity. Am. J. Physiol.-Leg. Content 46, 11–21. 10.1152/ajplegacy.1918.46.1.11.

37. Stewart, B.J., Ferdinand, J.R., Young, M.D., Mitchell, T.J., Loudon, K.W., Riding, A.M., Richoz, N., Frazer, G.L., Staniforth, J.U.L., Vieira Braga, F.A., et al. (2019). Spatiotemporal immune zonation of the human kidney. Science 365, 1461–1466. 10.1126/science.aat5031.

38. Weissenbacher, A., Huang, H., Surik, T., Lo Faro, M.L., Ploeg, R.J., Coussios, C.C., Friend, P.J., and Kessler, B.M. (2021). Urine recirculation prolongs normothermic kidney perfusion via more optimal metabolic homeostasis—a proteomics study. Am. J. Transplant. 21, 1740–1753. 10.1111/ajt.16334.

39. Mellati, A., Lo Faro, L., Dumbill, R., Meertens, P., Rozenberg, K., Shaheed, S., Snashall, C., McGivern, H., Ploeg, R., and Hunter, J. (2022). Kidney Normothermic Machine Perfusion Can Be Used as a Preservation Technique and a Model of Reperfusion to Deliver Novel Therapies and Assess Inflammation and Immune Activation. Front. Immunol. 13.

40. McEvoy, C.M., Clotet-Freixas, S., Tokar, T., Pastrello, C., Reid, S., Batruch, I., RaoPeters, A.A.E., Kaths, J.M., Urbanellis, P., Farkona, S., et al. (2021). Normothermic Ex-vivo Kidney Perfusion in a Porcine Auto-Transplantation Model Preserves the Expression of Key Mitochondrial Proteins: An Unbiased Proteomics Analysis. Mol. Cell. Proteomics MCP 20, 100101. 10.1016/j.mcpro.2021.100101.

41. Budhiraja, P., Reddy, K.S., Butterfield, R.J., Jadlowiec, C.C., Moss, A.A., Khamash, H.A., Kodali, L., Misra, S.S., and Heilman, R.L. (2022). Duration of delayed graft function and its impact on graft outcomes in deceased donor kidney transplantation. BMC Nephrol. 23, 154. 10.1186/s12882-022-02777-9.

42. Husen, P., Boffa, C., Jochmans, I., Krikke, C., Davies, L., Mazilescu, L., Brat, A., Knight, S., Wettstein, D., Cseprekal, O., et al. (2021). Oxygenated End-Hypothermic Machine Perfusion in Expanded Criteria Donor Kidney Transplant: A Randomized Clinical Trial. JAMA Surg. 156, 517–525. 10.1001/jamasurg.2021.0949.

43. Lindeman, J.H., Wijermars, L.G., Kostidis, S., Mayboroda, O.A., Harms, A.C., Hankemeier, T., Bierau, J., Sai Sankar Gupta, K.B., Giera, M., Reinders, M.E., et al. (2020). Results of an explorative clinical evaluation suggest immediate and persistent post-reperfusion metabolic paralysis drives kidney ischemia reperfusion injury. Kidney Int. 10.1016/j.kint.2020.07.026.

44. Bonventre, J.V., and Yang, L. (2011). Cellular pathophysiology of ischemic acute kidney injury. J. Clin. Invest. 121, 4210–4221. 10.1172/JCI45161.

45. Aukland, K., and Krog, J. (1960). Renal Oxygen Tension. Nature 188, 671–671. 10.1038/188671a0.

46. Fu, Y., Tang, C., Cai, J., Chen, G., Zhang, D., and Dong, Z. (2018). Rodent models of AKI-CKD transition. Am. J. Physiol.-Ren. Physiol. 315, F1098–F1106. 10.1152/ajprenal.00199.2018.

47. Vogel, C., and Marcotte, E.M. (2012). Insights into the regulation of protein abundance from proteomic and transcriptomic analyses. Nat. Rev. Genet. 13, 227–232. 10.1038/nrg3185.

48. de Sousa Abreu, R., Penalva, L.O., Marcotte, E.M., and Vogel, C. (2009). Global signatures of protein and mRNA expression levels. Mol. Biosyst. 5, 1512–1526. 10.1039/b908315d.

49. Molkentin, J.D., Bugg, D., Ghearing, N., Dorn, L.E., Kim, P., Sargent, M.A., Gunaje, J., Otsu, K., and Davis, J. (2017). Fibroblast-Specific Genetic Manipulation of p38 Mitogen-Activated Protein Kinase In Vivo Reveals Its Central Regulatory Role in Fibrosis. Circulation 136, 549–561. 10.1161/CIRCULATIONAHA.116.026238.

50. Wissing, E.R., Boyer, J.G., Kwong, J.Q., Sargent, M.A., Karch, J., McNally, E.M., Otsu, K., and Molkentin, J.D. (2014). P38α MAPK underlies muscular dystrophy and myofiber death through a Bax-dependent mechanism. Hum. Mol. Genet. 23, 5452–5463. 10.1093/hmg/ddu270.

51. Matsuoka, H., Arai, T., Mori, M., Goya, S., Kida, H., Morishita, H., Fujiwara, H., Tachibana, I., Osaki, T., and Hayashi, S. (2002). A p38 MAPK inhibitor, FR-167653, ameliorates murine bleomycin-induced pulmonary fibrosis. Am. J. Physiol.-Lung Cell. Mol. Physiol. 283, L103–L112. 10.1152/ajplung.00187.2001.

52. Stambe, C., Atkins, R.C., Tesch, G.H., Masaki, T., Schreiner, G.F., and Nikolic-Paterson, D.J. (2004). The Role of p38α Mitogen-Activated Protein Kinase Activation in Renal Fibrosis. J. Am. Soc. Nephrol. 15, 370. 10.1097/01.ASN.0000109669.23650.56.

53. di Mari, J.F., Davis, R., and Safirstein, R.L. (1999). MAPK activation determines renal epithelial cell survival during oxidative injury. Am. J. Physiol. 277, F195–203. 10.1152/ajprenal.1999.277.2.F195.

54. Park, K.M., Kramers, C., Vayssier-Taussat, M., Chen, A., and Bonventre, J.V. (2002). Prevention of Kidney Ischemia/Reperfusion-induced Functional Injury, MAPK and MAPK Kinase Activation, and Inflammation by Remote Transient Ureteral Obstruction*. J. Biol. Chem. 277, 2040–2049. 10.1074/jbc.M107525200.

55. Kwon, D.S., Kwon, C.H., Kim, J.H., Woo, J.S., Jung, J.S., and Kim, Y.K. (2006). Signal transduction of MEK/ERK and PI3K/Akt activation by hypoxia/reoxygenation in renal epithelial cells. Eur. J. Cell Biol. 85, 1189–1199. 10.1016/j.ejcb.2006.06.001.

56. Yin, T., Sandhu, G., Wolfgang, C.D., Burrier, A., Webb, R.L., Rigel, D.F., Hai, T., and Whelan, J. (1997). Tissue-specific Pattern of Stress Kinase Activation in Ischemic/Reperfused Heart and Kidney *. J. Biol. Chem. 272, 19943–19950. 10.1074/jbc.272.32.19943.

57. Liu, D., Liu, X., Su, Y., and Zhang, X. (2011). Renal expression of proto-oncogene Ets-1 on matrix remodeling in experimental diabetic nephropathy. Acta Histochem. 113, 527– 533. 10.1016/j.acthis.2010.05.006.

58. Naito, T., Tanihata, Y., Nishimura, H., Tanaka, T., Higuchi, C., Taguchi, T., and Sanaka, T. (2005). Expression of matrix metalloproteinase-9 associated with ets-1 proto-oncogene in rat tubulointerstitial cells. Nephrol. Dial. Transplant. 20, 2333–2348. 10.1093/ndt/gfi013.

59. Hosgood, S.A., Heurn, E. van, and Nicholson, M.L. (2015). Normothermic machine perfusion of the kidney: better conditioning and repair? Transpl. Int. 28, 657–664. 10.1111/tri.12319.

60. Ogurlu, B., Pamplona, C.C., Van Tricht, I.M., Hamelink, T.L., Lantinga, V.A., Leuvenink, H.G.D., Moers, C., and Pool, M.B.F. (2023). Prolonged Controlled Oxygenated Rewarming Improves Immediate Tubular Function and Energetic Recovery of Porcine Kidneys During Normothermic Machine Perfusion. Transplantation 107, 639–647. 10.1097/TP.0000000000004427.

61. Dumbill, R., Rabcuka, J., Fallon, J., Knight, S., Hunter, J., Voyce, D., Barrett, J.T., Ellen, M., Weissenbacher, A., Kurniawan, T., et al. (2023). Impaired O2 unloading from stored blood results in diffusion-limited O2 release at tissues: evidence from human kidneys. Blood, blood.2023022385. 10.1182/blood.2023022385.

62. Ferdinandy, P., Schulz, R., and Baxter, G.F. (2007). Interaction of Cardiovascular Risk Factors with Myocardial Ischemia/Reperfusion Injury, Preconditioning, and Postconditioning. Pharmacol. Rev. 59, 418–458. 10.1124/pr.107.06002.

63. Ferdinandy, P., Szilvassy, Z., and Baxter, G.F. (1998). Adaptation to myocardial stress in disease states: is preconditioning a healthy heart phenomenon? Trends Pharmacol. Sci. 19, 223–229. 10.1016/S0165-6147(98)01212-7.

64. Lohmann, S., Eijken, M., Møldrup, U., Møller, B.K., Hunter, J., Moers, C., Ploeg, R.J., Baan, C.C., Jespersen, B., and Keller, A.K. (2019). A Pilot Study of Postoperative Animal Welfare as a Guidance Tool in the Development of a Kidney Autotransplantation Model With Extended Warm Ischemia. Transplant. Direct 5, e495. 10.1097/TXD.0000000000000941.

65. Shaheed, S., Rustogi, N., Scally, A., Wilson, J., Thygesen, H., Loizidou, M.A., Hadjisavvas, A., Hanby, A., Speirs, V., Loadman, P., et al. (2013). Identification of Stage-Specific Breast Markers Using Quantitative Proteomics. J. Proteome Res. 12, 5696–5708. 10.1021/pr400662k.

66. Ahlmann-Eltze, C., and Anders, S. proDA: Probabilistic Dropout Analysis for Identifying Differentially Abundant Proteins in Label-Free Mass Spectrometry. 26.

67. Ritchie, M.E., Phipson, B., Wu, D., Hu, Y., Law, C.W., Shi, W., and Smyth, G.K. (2015). limma powers differential expression analyses for RNA-sequencing and microarray studies. Nucleic Acids Res. 43, e47. 10.1093/nar/gkv007.

68. Johnson, W.E., Li, C., and Rabinovic, A. (2007). Adjusting batch effects in microarray expression data using empirical Bayes methods. Biostat. Oxf. Engl. 8, 118–127. 10.1093/biostatistics/kxj037.

69. The UniProt Consortium (2023). UniProt: the Universal Protein Knowledgebase in 2023. Nucleic Acids Res. 51, D523–D531. 10.1093/nar/gkac1052.

70. The Gene Ontology Consortium, Carbon, S., Douglass, E., Good, B.M., Unni, D.R., Harris, N.L., Mungall, C.J., Basu, S., Chisholm, R.L., Dodson, R.J., et al. (2021). The Gene Ontology resource: enriching a GOld mine. Nucleic Acids Res. 49, D325–D334. 10.1093/nar/gkaa1113.

71. Korotkevich, G., Sukhov, V., Budin, N., Shpak, B., Artyomov, M.N., and Sergushichev, A. (2021). Fast gene set enrichment analysis. Preprint at bioRxiv, 10.1101/060012 https://doi.org/10.1101/060012.

72. Garcia-Alonso, L., Iorio, F., Matchan, A., Fonseca, N., Jaaks, P., Peat, G., Pignatelli, M., Falcone, F., Benes, C.H., Dunham, I., et al. (2018). Transcription Factor Activities Enhance Markers of Drug Sensitivity in Cancer. Cancer Res. 78, 769–780. 10.1158/0008-5472.CAN-17-1679.

73. Badia-i-Mompel, P., Vélez Santiago, J., Braunger, J., Geiss, C., Dimitrov, D., Müller-Dott, S., Taus, P., Dugourd, A., Holland, C.H.,Ramirez Flores, R.O., et al. (2022). decoupleR: ensemble of computational methods to infer biological activities from omics data. Bioinforma. Adv. 2, vbac016. 10.1093/bioadv/vbac016.

74. Lake, B.B., Menon, R., Winfree, S., Hu, Q., Ferreira, R.M., Kalhor, K., Barwinska, D., Otto, E.A., Ferkowicz, M., Diep, D., et al. (2021). An atlas of healthy and injured cell states and niches in the human kidney. Preprint at bioRxiv, 10.1101/2021.07.28.454201 https://doi.org/10.1101/2021.07.28.454201.

75. Hao, Y., Hao, S., Andersen-Nissen, E., Mauck, W.M., Zheng, S., Butler, A., Lee, M.J., Wilk, A.J., Darby, C., Zager, M., et al. (2021). Integrated analysis of multimodal single-cell data. Cell 184, 3573–3587.e29. 10.1016/j.cell.2021.04.048.

76. Subramanian, A., Tamayo, P., Mootha, V.K., Mukherjee, S., Ebert, B.L., Gillette, M.A., Paulovich, A., Pomeroy, S.L., Golub, T.R., Lander, E.S., et al. (2005). Gene set enrichment analysis: A knowledge-based approach for interpreting genome-wide expression profiles. Proc. Natl. Acad. Sci. 102, 15545–15550. 10.1073/pnas.0506580102.

