## Supplementary Information for "End of preservation normothermic machine perfusion of porcine kidneys after ischaemic injury reprograms metabolism and induces fibrosis after transplant despite unchanged function: insights from the renal proteome"

### Supporting information

#### 1. Methods

##### 1.1 Porcine autotransplantation model

The left kidney was procured after being subjected to 75 min of warm ischaemia: previously demonstrated in our hands to induce severe ischaemic injury whilst maintaining acceptable animal welfare[64]. The graft was flushed with 500 mL of cold Belzer UW Cold Storage Solution (UW-CSS; Bridge to Life) and then preserved according to the experimental group. Kidneys that had undergone NMP were flushed and cooled with 500 mL UW-CSS. The right kidney was then removed and the left kidney autotransplanted to the right renal artery, vein and ureter using an end-to-end anastomosis. Healthy control kidneys were obtained from pigs that had been subject to a non-invasive infusion of 25 mL phosphate buffered saline as a placebo procedure.

##### 1.2 *Ex vivo* kidney perfusion

For both oxHMP and NMP, perfusion was performed as reported previously[24]. A custom perfusion circuit consisted of an oxygenator (Terumo, Denmark), a centrifugal pump delivering pulsatile perfusion (Medos Deltastream DP2 and Pumpdrive DP2, Medos Medizintechnik AG), an organ chamber with a 3 mm straight cannula (Organ Recovery Systems), and pressure (TruWave disposable pressure transducer, Edwards Lifesciences) and flow (Transonic Systems Europe BV) probes. Any urine produced was recirculated into the perfusate reservoir.

Oxygenated HMP was performed for 14 hours at a temperature of 4-7 °C and a mean arterial pressure of 25 mmHg, using 500 mL of University of Wisconsin machine perfusion solution (UW-MPS; Bridge to Life Ltd) which was actively oxygenated with 0.1 L/min oxygen.

Before commencing NMP, kidneys were first flushed with 50 ml cold Ringer's acetate (Fresenius Kabi). NMP was then performed for 4 hours at 37 °C and a mean arterial pressure of 70 mmHg. The perfusate consisted of 170 ml allogeneic porcine erythrocytes diluted in of 250 mL 5% (w/v) human albumin (Alburex, CSL Behring) to give haematocrit of 25–30%. This was supplemented with 5 ml sodium bicarbonate 8.4% (w/v; B. Braun), 6 ml glucose 5% (w/v; B. Braun), 5 IU insulin (Novo Nordisk A/S), 3 ml calcium gluconate 10% (w/v; B. Braun), 10 mg mannitol (Sigma-Aldrich, UK), and creatinine to a concentration of 1000 µM (SigmaAldrich, UK). An initial bolus of 300 mg Amoxicillin-clavulanate (Bowmed) was followed by a further 60 mg every hour. Similarly, an initial bolus dose of 1.25 mg verapamil (Orion Pharma) was followed by a continuous infusion at 0.25 mg/h.

##### 1.3 Measurement of kidney function

On day 14 after transplantation, glomerular filtration rate was measured (mGFR) by the clearance of 99mTc-diethylenetriaminepentaacetic acid (DTPA) with a priming dose of 75 MBq and then infusion at 37.5 MBq/h as previously described (Lohmann et al., 2021). Following measurement of kidney function, the graft was removed and the pigs euthanised.

#### 1.4 Protein extraction, digestion and sample multiplexing for mass spectrometry

To extract proteins RIPA lysis and extraction buffer (Sigma Aldrich, UK) supplemented with 1% protease inhibitor (Thermo Fisher Scientific, UK) was added to the samples at 3 times the weight of biopsy material. After 3 cycles of probe sonication for 3 seconds on ice, the mixture was centrifuged at 4 °C for 20 min at 13000 rpm. The pelleted cellular debris were discarded and the supernatant was transferred to a new centrifuge tube. Protein concentration was determined by BCA assay (Thermo Fisher Scientific, UK). 10 µg of protein per sample was separated by SDS-PAGE and stained by Coomasie brilliant blue to visually verify sample quality.

Approximately 100 µg of protein from each sample was precipitated using ice-cold acetone. The precipitated proteins were dissolved in 0.1 M Triethylammonium bicarbonate (TEAB) with 1% w/v SDS, reduced and alkylated with 0.5 M TCEP for 1 hour at 37 °C and 10 mM iodoacetamide for 30 min at room temperature and in the dark, respectively. The protein solution was diluted 10-fold using 0.1 M TEAB and Pierce Trypsin Protease, MS Grade (Thermo Fisher Scientific, UK) was added at 1:50 mass ratio (trypsin:protein) at 37 °C for 4 hours and followed by a second step of trypsin at a mass ratio of 1:100 (trypsin:protein) overnight. Trypsin enzymatic peptides were dried under vacuum after desalination by SPE Cartridge Bond Elut C18 columns (Agilent Technologies, Santa Clara, USA). The peptides were dissolved with 0.5 M TEAB (Sigma Aldrich, UK) and labelled with isobaric mass tags according to the manufacturer instructions of the 10plex TMT kit (Thermo Fisher Scientific, UK). Samples were allocated randomly between TMT channels and batches. A pool sample was used to control batch effects between sets of TMT labelling.

#### 1.5 LC-MS/MS data processing and database searching

Proteome Discover (version 2.2, Thermo Fisher Scientific, UK), was used for identification and quantification of proteins using an in-house Sequest HT database search program (University of Washington, Seattle, Washington, USA), utilising the Sus scrofa database obtained from Uni-Prot (version 2021 containing 119,328 protein sequences). Data processing parameters were follows: enzymatic digestion by trypsin with up to 2 missed cleavage sites, a 10 ppm mass error of primary parent ions, a 0.05 Da mass error of secondary fragment ions, cysteine alkylation as a fixed modification and methionine oxidation as a variable modification. The quantitative method was set as TMT 10plex; protein identification by peptide spectrum matching identification was done at a false discovery rate of 1%. Only proteins containing at least one unique peptide and ≥2 PSMs were retained for further analysis.

#### 1.6 Western blot analysis

Western blot experiments were performed as described previously[65]. Briefly, equal amounts of proteins were separated using a 12% SDS-poly-acrylamide gel, transferred by electrophoresis to nitrocellulose membranes and blocked with 5% milk. Membranes were washed and probed with corresponding primary antibodies and subsequently by secondary antibodies. Protein bands were visualised by LI-COR Odyssey® DLx Imaging System (LI-COR Biosciences, Ltd. UK). Gel Analyzer 19.1 (http://www.gelanalyzer.com/), was used to analyse the images and quantify bands. The intensity of protein bands was normalised to β-actin intensity and a rolling ball algorithm was applied to remove the background intensity in case of uneven exposure. Antibodies used were as follows:

Antidbodies used were as follows: mouse monoclonal anti-COL1A1 (dilution 1:1,000; Cat#67288-1-Ig, Proteintech, Rosemont, IL, USA), Rabbit polyclonal to beta Actin (dilution 1:2,000, Cat#ab8227) from Abcam PLC, Cambridge UK, IRDye® 680RD Goat anti-mouse

IgG secondary antibody (dilution 1:2,000; Cat#926-68070) and IRDye® 800CW Goat anti-rabbit IgG secondary antibody (dilution 1:2,000; Cat#926-32211) from LI-COR Biosciences, Ltd. UK.

##### 1.7 Statistical Analysis

Within each set of TMT reagents, data were expressed relative to the pool channel. A variance stabilising transformation was used to account for mean-variance bias. Missing values were accounted for using a Bayesian probabilistic dropout model[66] to account for the fact that missingness is dependent upon peptide abundance. To calculate the differential abundance of proteins between experimental conditions, an empirical bayes moderated t-statistic was calculated with the batch specified as a covariate[67]. For downstream analysis and visualisation, batch correction was instead performed using the ComBat algorithm[68].

Porcine proteins were mapped to human genes using the UniProt[69] mapping tool. Gene set enrichment analysis of the Gene Ontology resource[70] biological process gene sets was performed with the fgsea algorithm[71] using as the rank metric the $\log_2$ fold change between experimental conditions. Dorothea[72] was used to calculate enriched transcriptional regulators and decoupleR[73] to calculate pathway enrichment between conditions, using the consensus of the weighted mean and univariate linear model statistics.

To identify cell types that are changing, we intersect our proteomics data with a single cell RNAseq dataset previously generated by Stewart et al[37] and now available via the kidney cell atlas. Cell types were annotated by integration with an annotated reference dataset[74] using Azimuth[75]. Markers for the "annotation.l1" celltypes were defined using the Wilcoxon signed rank test implemented in Seurat[75]. Enrichment of cell types defined by these markers was performed using the fgsea algorithm[71], to approximate the method of Subramanian and colleagues[76].

#### 2. Results

##### 2.1 Sample quality assurance

After protein extraction, the concentration of protein in samples ranged from 5.2 µg/µl to 10.2 µg/µl, providing enough material for downstream analysis in all cases. Approximately 20 µg of protein from each sample was separated using sodium dodecyl-sulfate polyacrylamide gel electrophoresis (SDS-PAGE) and stained with Coomassie brilliant blue. This showed clear and uniform bands, which visually confirm the quality of the samples and the extraction procedure (Fig S2A).

##### 2.2 Mass Spectrometry

In this project, 335,625 MS/MS were acquired by mass spectrometry. After searching the Uniprot (protein sequence database), 114,386 valid PSM (peptide spectral match) were obtained, and the utilization rate of the spectrograms was 34.1%. A total of 2,848 peptides were identified by spectrographic analysis, of which there were 2,418 unique peptides. Among the identified peptides, 75% of peptides had a mass error less then ±5 ppm, while 25% peptides had mass errors from ±5 ppm to ±10ppm. These results showed an acceptable mass accuracy of the mass spectrometry data. The coverage of all proteins ranged from 1% to 68%; the number of proteins with a coverage greater than 5% accounted for 60% of identified proteins. The length of the majority of the peptides (83%) ranged from 7 to 21 amino acids which confirmed the complete digestion of the proteins in our sample preparation.
