## Supplementary Figures for "End of preservation normothermic machine perfusion of porcine kidneys after ischaemic injury reprograms metabolism and induces fibrosis after transplant despite unchanged function: insights from the renal proteome"

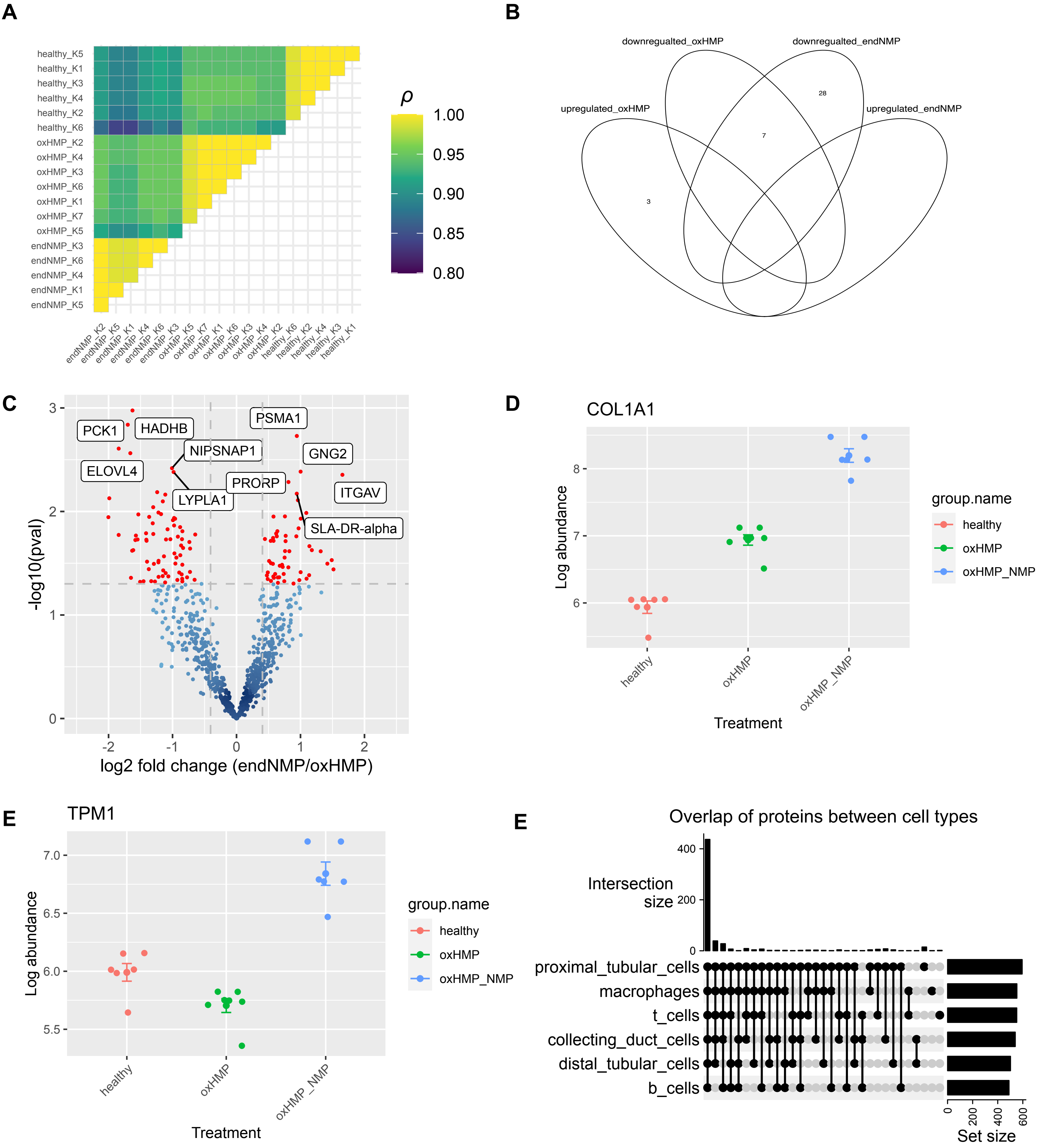


**Figure S1. A.** Pearson's rho between all combinations of samples, measured across all proteins, shows that samples within an experimental group are more similar to each other than to other samples. **B.** Overlap of significantly regulated gene sets between either oxHMP or endNMP kidneys and healthy controls. **C.** Protein abundance changes between endNMP and oxHMP **D.** Abundance of COL1A1 measured by LC-MSMS E. Abundance of TPM1 measured by LC-MSMS F. Overlap of gene expression between cell types.

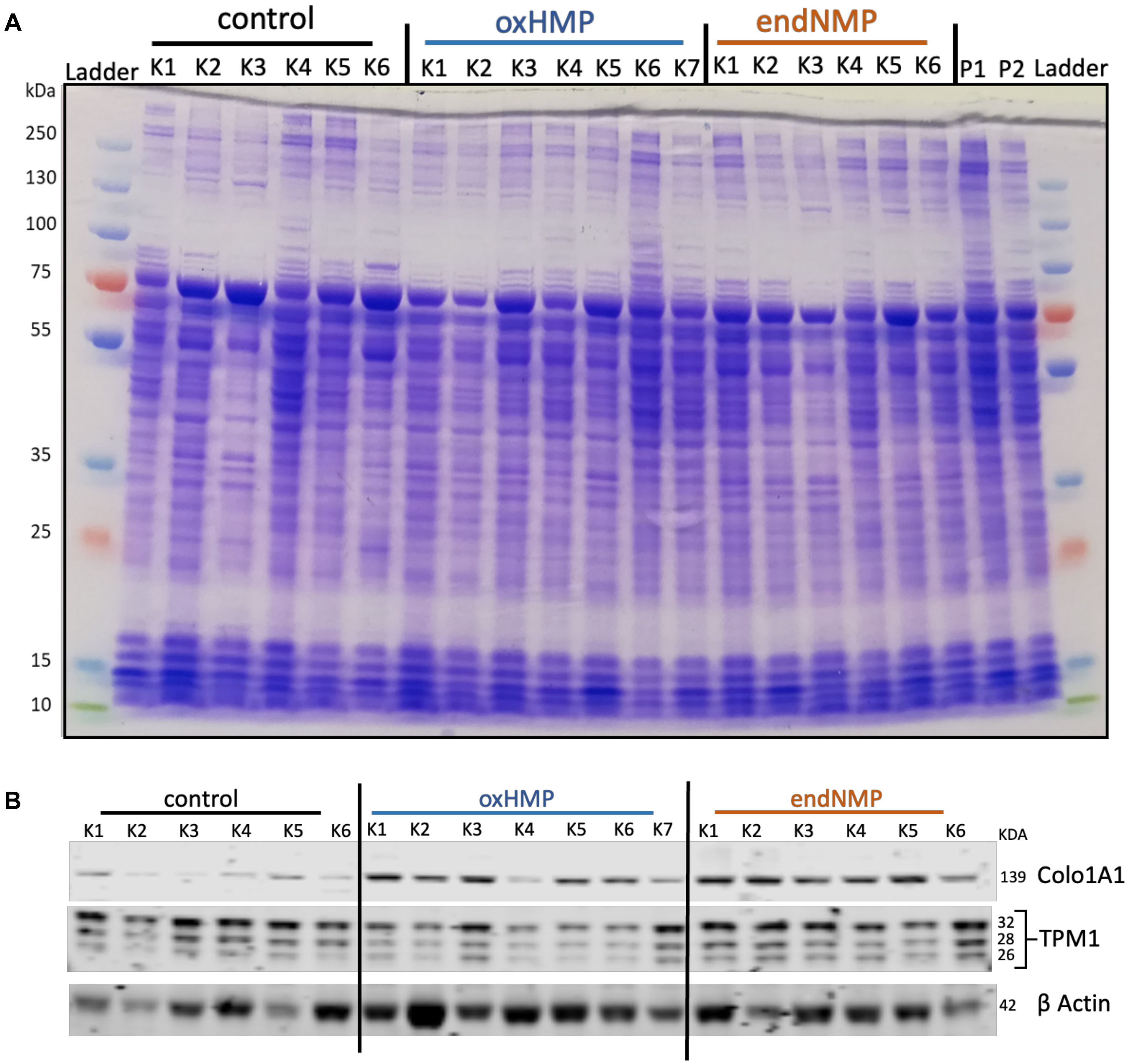


**Figure S2. A.** SDS-PAGE analysis of kidney biopsy total proteins. 20μg of extracted proteins from each sample was separated by molecular weight using sodium dodecyl sulphate-polyacrylamide gel electrophoresis (SDS-PAGE) and stained with Coomassie blue dye. **B.** Image of western blot.

K - kidney, P - pooled sample
